# A virus-encoded microRNA contributes to evade innate immune response during SARS-CoV-2 infection

**DOI:** 10.1101/2021.09.09.459577

**Authors:** Meetali Singh, Maxime Chazal, Piergiuseppe Quarato, Loan Bourdon, Christophe Malabat, Thomas Vallet, Marco Vignuzzi, Sylvie van der Werf, Sylvie Behillil, Flora Donati, Nathalie Sauvonnet, Giulia Nigro, Maryline Bourgine, Nolwenn Jouvenet, Germano Cecere

## Abstract

SARS-CoV-2 infection results in impaired interferon response in severe COVID-19 patients. However, how SARS-CoV-2 interferes with host immune response is incompletely understood. Here, we sequenced small RNAs from SARS-CoV-2-infected human cells and identified a micro-RNA (miRNA) encoded in a recently evolved region of the viral genome. We show that the virus-encoded miRNA produces two miRNA isoforms in infected cells by the enzyme Dicer and they are loaded into Argonaute proteins. Moreover, the predominant miRNA isoform targets the 3’UTR of interferon-stimulated genes and represses their expression in a miRNA-like fashion. Finally, the two viral miRNA isoforms were detected in nasopharyngeal swabs from COVID-19 patients. We propose that SARS-CoV-2 employs a virus-encoded miRNA to hijack the host miRNA machinery and evade the interferon-mediated immune response.

## INTRODUCTION

The infection by the severe acute respiratory syndrome-related coronavirus 2 (SARS-CoV-2), which causes Coronavirus Disease-2019 (COVID-19), is characterized by a wide range of symptoms, which in some cases lead to severe or critical disease outcome, including pneumonia and acute respiratory failure (Huang *et al*, 2020; Salje *et al*, 2020). Several studies have highlighted the central role of interferons (IFNs) in the outcome of COVID-19 disease (Acharya *et al*, 2020; Kim & Shin, 2021; Schultze & Aschenbrenner, 2021). The production of IFNs results in the activation of hundreds of interferon-stimulated genes (ISGs), which are the effectors of the host innate antiviral response (Lazear *et al*, 2019). However, patients with severe and critical COVID-19 disease manifestation show an impaired type I IFN response (Hadjadj *et al*, 2020). Moreover, several human cell lines, primary cells and *in vivo* samples derived from COVID-19 patients display a general impairment in the activation of ISGs upon SARS-CoV-2 infection (Blanco-Melo *et al*, 2020; Zhang *et al*, 2020a; Bastard *et al*, 2020; Galani *et al*, 2021). Different SARS-CoV-2 encoded proteins have now been shown to interfere with the interferon response (Xia *et al*, 2020; Schroeder *et al*, 2021; Lei *et al*, 2020; Miorin *et al*, 2020; Lin *et al*, 2021; Wu *et al*, 2021), implicating the importance on type I IFN response for counteracting SARS-CoV-2 infection.

Among the different mechanisms employed by viruses to interfere with host innate immune responses includes the use of small regulatory RNAs. Small RNAs, such as microRNAs (miRNAs), are fundamental regulators of host gene expression programs, including antiviral innate immunity genes (Girardi *et al*, 2018). They are encoded by the host genome in regions that form stem-loop RNA structures (Kim *et al*, 2009), which are processed by the endoribonuclease enzyme Dicer resulting in small RNAs of approximately 22 nucleotides (nt) in length. The mature miRNA is then loaded by Argonaute proteins (AGOs), which are a part of the RNA silencing effector complex that regulates target transcripts by sequence complementarity (Bartel, 2018). Dicer also cleaves doublestranded RNAs (dsRNAs) derived from RNA viruses’ genomes to inhibit viral replication and induce viral immunity through the production of small interfering RNAs (siRNAs) (Berkhout, 2018). Moreover, miRNAs can also be encoded by viral genomes and processed by the host miRNA pathway (Mishra *et al*, 2020). The functions of virus-encoded miRNAs are still not fully understood. However, in some cases, the virus can employ miRNAs to evade host immune response (Mishra *et al*, 2020). Given that some of the host enzymes involved in the biogenesis of miRNAs localize to the nucleus (Kim *et al*, 2009), all known viral miRNAs are encoded by RNA and DNA viruses replicating into the nucleus, such as Herpesviruses and Influenza virus (Mishra *et al*, 2020). Nonetheless, cytoplasmic RNA viruses carrying artificial miRNA sequences can produce mature miRNAs (Rouha *et al*, 2010; Shapiro *et al*, 2012; Langlois *et al*, 2012). Therefore, whether cytoplasmic RNA viruses, such as SARS-CoV-2, also produce their own miRNAs remains to be investigated.

## RESULTS

To analyze the repertoire of small RNAs produced upon SARS-CoV-2 infection with potential regulatory functions, we generated 5’ monophosphate-dependent small RNA libraries from human colorectal adenocarcinoma cells (Caco-2), an intestinal cellular model and human pulmonary ACE2-expressing A549 (A549-ACE2) cells, both known to be highly susceptible to SARS-CoV-2 infection (Takayama, 2020; Chu *et al*, 2020). Specifically, we gel-purified and sequenced small RNAs, ranging from 18 to 26 nucleotides (nt), at 24- and 48-hours post-infection (hpi) together with their respective noninfected control cells. Our analysis revealed the presence of reads mapping to the SARS-CoV-2 genome in infected versus non-infected cells (Table S1). Their amount increased over the course of infection, indicating that these reads are produced during viral replication in both cellular models tested (Fig. S1A-B and S2A-B). Next, we analyzed their size distribution and directionality to verify whether these small RNAs are generated by Dicer from the viral dsRNA replication intermediates. We found a very small fraction of SARS-CoV-2 reads mapping in antisense orientation and the majority of reads were not enriched for 22 nt reads (Fig. S1C and S2C-D), the typical size of Dicer-cleaved small RNAs (Bernstein *et al*, 2001). In contrast, the small RNAs mapped to the human genome, which included a large fraction of known human miRNAs (Table S2), were enriched for small RNAs of 22 nt in length (Fig. S1D and S2E). These results suggest that the majority of the SARS-CoV-2 reads do not represent canonical siRNAs generated by Dicer from dsRNA intermediates but instead result from degradation fragments of the viral RNA genome. Accordingly, most of the SARS-CoV-2 reads were distributed across the whole length of the viral genome (Fig. S1A and S2A).

Intriguingly, we identified a well-defined small RNA peak mapping to the beginning of the ORF7a, which increased in abundance upon viral replication (Fig. S1A and S2A). The analysis of the size distribution of the reads across 200 nt surrounding the identified peak, showed an enrichment of reads of 22 nt in length (Fig. S1E and S2F). This result suggests that the small RNAs derived from this region do not result from the viral genome’s degradation and could instead derive from virus-encoded miRNAs. To identify the precise sequences of these 22 nt small RNAs, we analyzed all the 22 nt reads that mapped to the SARS-CoV-2 genome (Fig. 1A). Our analysis showed the presence of two predominant 22 nt small RNA sequences, which differ in only 2 nt and correspond to the identified peak at the beginning of the ORF7a in both Caco-2 and A549-ACE2 cells (Fig. 1B-C). These results indicate that the two predominant 22 nt small RNAs derived from the ORF7a are produced in intestinal- and pulmonary-derived human cell lines, which represent tissues targeted by SARS-CoV-2 in humans (Wu *et al*, 2020). In addition to these small RNAs, we identified another abundant 22 nt small RNA derived from ORF1b in A549-ACE2 infected cells (Fig. 1A,C). However, this small RNA is present at a much lower level in Caco-2 cells (Fig. 1A-B), and we thus decided to focus on the two small RNAs derived from the ORF7a.

**Figure 1:**
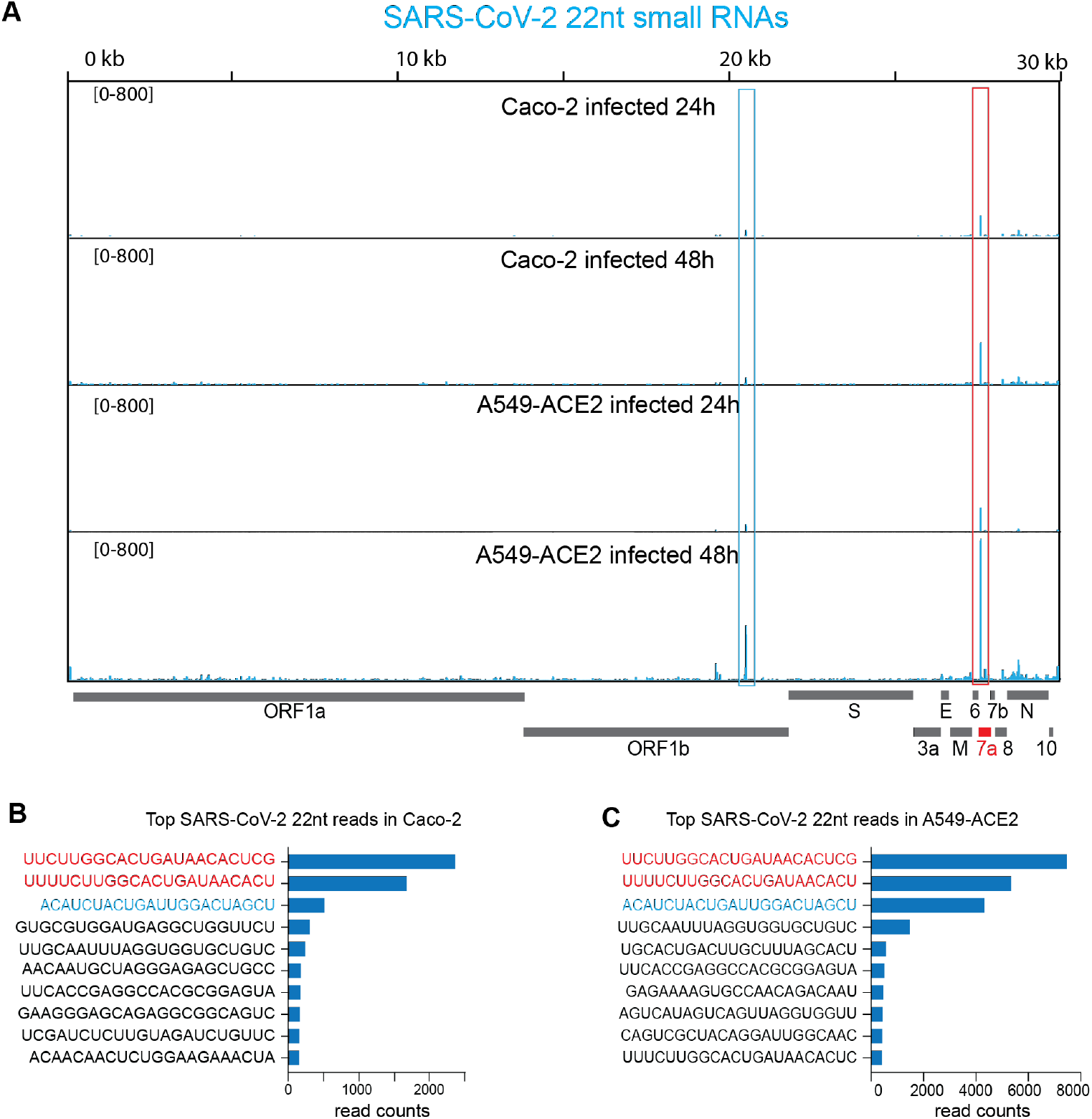
Identification of a virus-encoded miRNA during SARS-CoV-2 infection. **(A)** SARS-CoV-2 genomic view showing the distribution of normalized 22nt small RNA reads from Caco-2 and A549-ACE2 cells at 24 and 48 hpi. The most abundant small RNAs are marked by the red and blue boxes. For all the experiments shown n=2. **(B)** Average read counts for the ten most abundant 22 nt SARS-CoV-2 small RNAs in Caco-2 cells at 48 hpi. The two most abundant small RNAs which differ by only 2 nt, marked in red, are encoded from the ORF7a region marked by the red box in (A). **(C)** Average read counts for the ten most abundant 22 nt SARS-CoV-2 small RNAs in A549-ACE2 cells at 48 hpi. The two most abundant small RNAs encoded from the ORF7a region marked by the red box in (A) are marked in red. The third abundant small RNA, marked in blue, derived from the ORF1b region marked by the blue box in (A).

To verify that the two small RNAs can be viral-encoded miRNAs derived from a common stem-loop RNA precursor we analyzed the first 70 nt of the ORF7a containing the two small RNAs. Indeed, our analysis predicts the formation of a stem-loop structure (Fig. 2A), which could be recognized by Dicer to produce miRNAs (Lee *et al*, 2003). Thus, the two small RNAs generated from ORF7a are possibly two isoforms of the same virus-encoded miRNA, which are usually generated in mammals by imprecise Dicer cleavage of the same miRNA precursor (Chiang *et al*, 2010). Thus, we named these two small RNAs CoV2-miR-7a.1 and CoV2-miR-7a.2.

**Figure 2:**
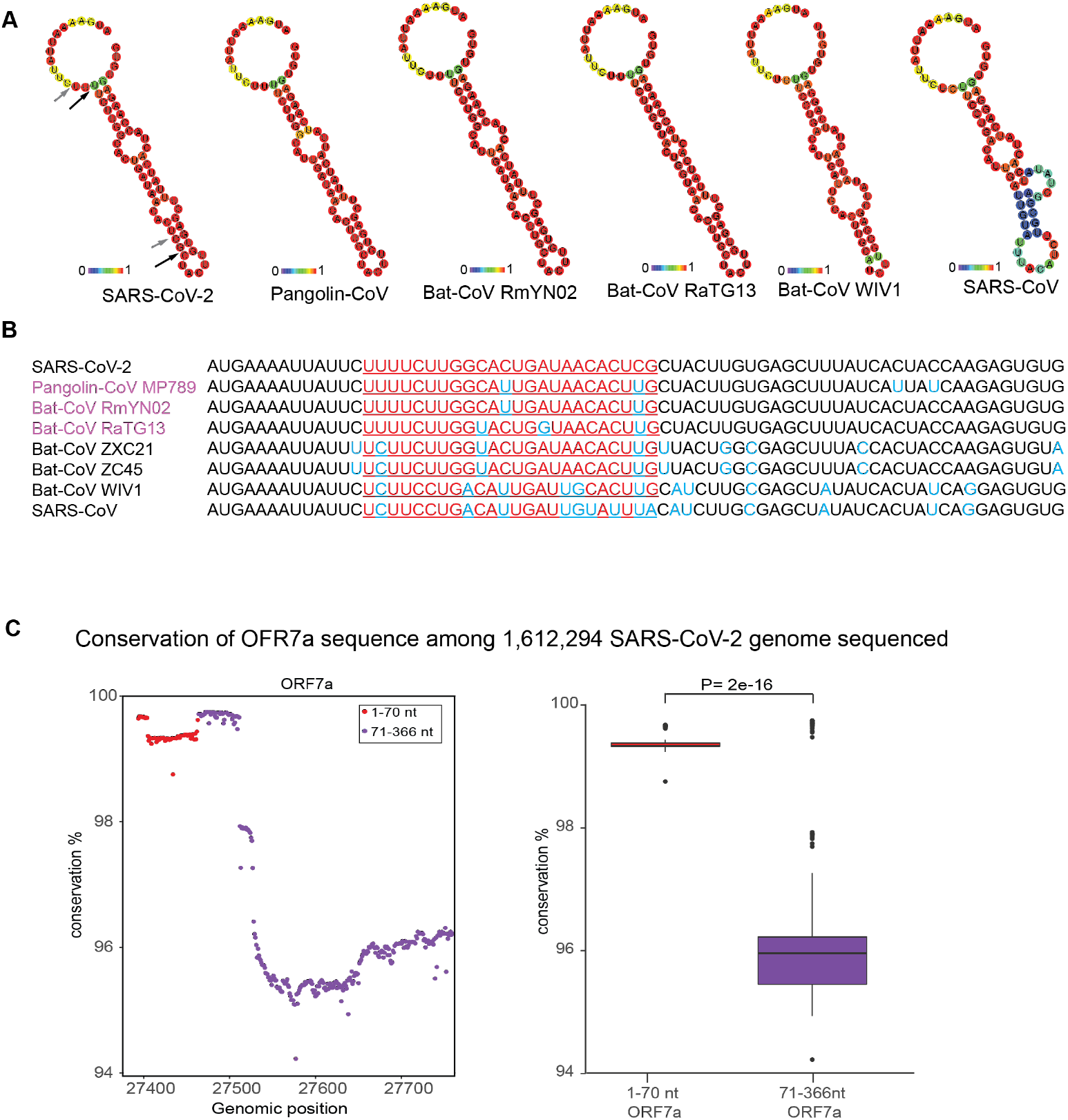
RNA secondary structure of SARS-CoV-2 miR-7a precursor and sequence conservation among different SARS coronaviruses and within SARS-CoV-2 variants. **(A)** Predicted RNA secondary structure for the CoV2-miR-7a and flanking sequence using the first 70 nt of the open reading frame of the ORF7a. The arrows indicate the sites of the miRNAs possibly cleaved by Dicer. The stem-loop structure is not conserved in SARS-CoV. The colors indicate the basepair probabilities. **(B)** Conservation of the first 70 nt of the ORF7a sequence among different SARS coronaviruses. The underlined sequences are related to the position of the SARS-CoV-2 miR-7a. The conserved ribonucleotides of the CoV2-miR-7a sequence are marked in red and in blue all the nonconserved ribonucleotides across the 70nt sequence. The bat and pangolin coronaviruses closely related to SARS-CoV-2 are marked in purple. (**C**) Percentage of conservation along the nucleotide positions in the ORF7a among 1,612,294 sequenced SARS-CoV-2 genomes. The First 70 nt are shown in red and shows a higher percentage of conservation compared to the rest of the sequence of ORF-7a. Boxplot shows the distribution of conservation percentage for each nucleotide either in the first 70 nt or 71-366 nt of ORF-7a among 1,612,294 sequenced SARS-CoV-2 genomes. Box plots display median (line), first and third quartiles (box), and 5th /95th percentile value (whiskers). Each dot represents the outliers. Two-tailed P values were calculated using the student’s t-test.

Interestingly, the sequence producing the CoV2-miR-7a is largely different in SARS-CoV genome (Fig. 2B), which does not generate the same stem-loop structure compatible with miRNA processing (Fig. 2A). Instead, the bat RmYN02 and the pangolin MP789/2019 coronaviruses, which are closely related to SARS-CoV-2 (Zhang *et al*, 2020b; Zhou *et al*, 2020a, 2020b), showed a similar sequence and predicted stem-loop structure unlike SARS-CoV and other bat coronaviruses, suggesting the recent evolution of the CoV2-miR-7a (Fig. 2A-B).

Next, we analyzed the degree of conservation within the ORF7a sequence among 1,612,294 genomic sequences of SARS-CoV-2 variants. We found that the first 70 nt encoding virus miRNAs are more conserved compared to the rest of the ORF7a among all the sequences of SARS-CoV-2 variants analyzed (Fig. 2C and Fig. 3). This result suggests that the beginning of the ORF7a is under selective pressure to maintain the sequence encoding the two miRNAs, which might be therefore biologically functional.

**Figure 3:**
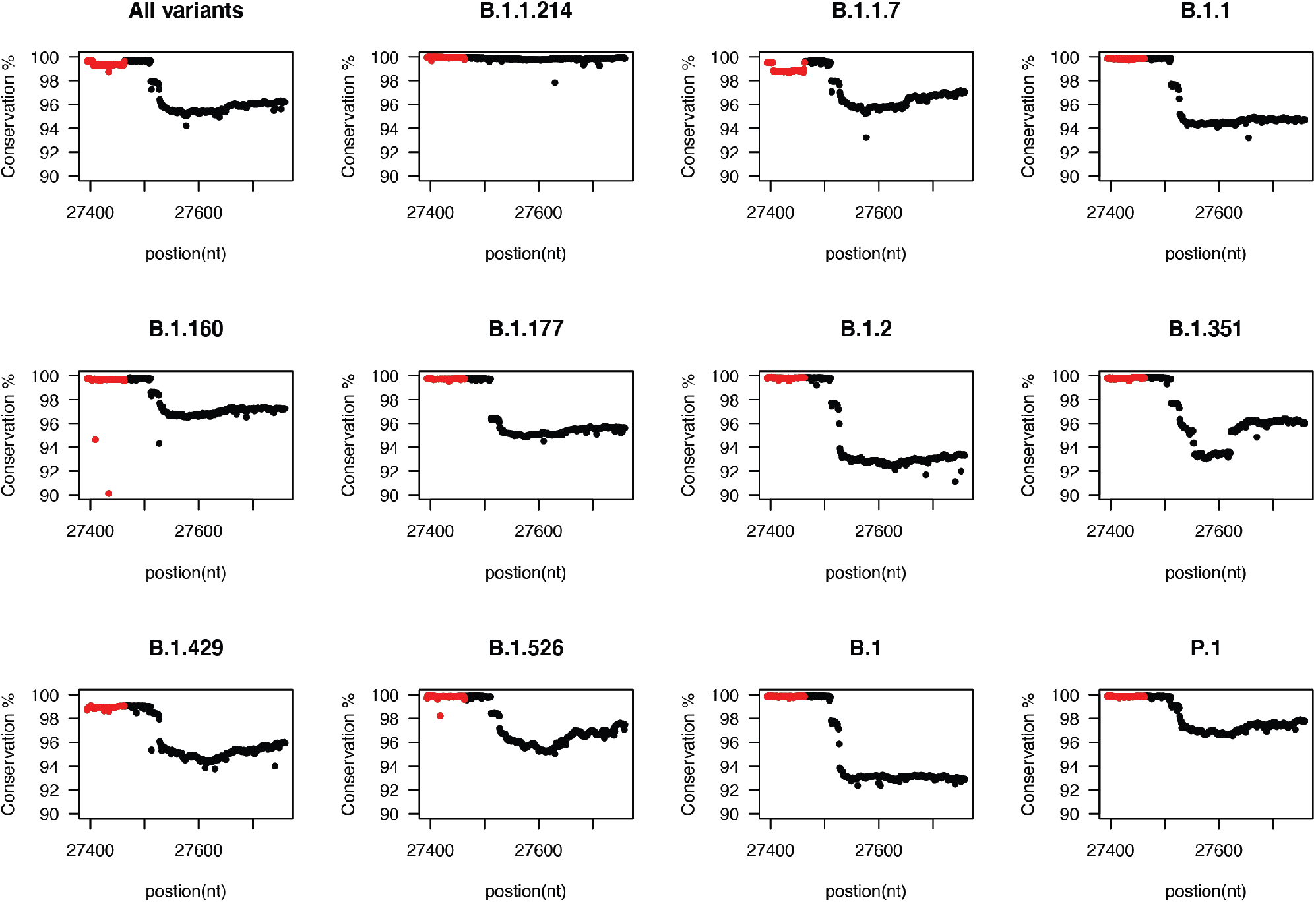
Conservation of first 70 nt and rest of the ORF-7a among different variants of SARS-CoV-2. Percentage of conservation along the nucleotide positions in the ORF7a among different variants of SARS-CoV-2. The First 70 nt are shown in red and shows a higher percentage of conservation compared to the rest of the sequence of ORF-7a. The number of genome sequences analyzed for each variant can be found in materials and methods

To test whether the human Dicer enzyme, DICER1, generates the CoV2-miR-7a.1 and CoV2-miR-7a.2, we used validated siRNAs to knock down DICER1 prior to SARS-CoV-2 infection in A549-ACE2 cells. We first validated the production of the CoV2-miR-7a. 1 and CoV2-miR-7a.2 by a stem-loop RT-qPCR assay commonly used to detect mature miRNAs (Chen *et al*, 2005). We were able to specifically detect the CoV2-miR-7a. 1 and CoV2-miR-7a.2 in A549-ACE2 infected cells, whereas no signal was obtained from a proximal region corresponding to the ORF6, which does not produce small RNAs (Fig. S3A). Moreover, we validated the sequencing results obtained in Caco-2 and A549-ACE2 cell lines showing that the CoV2-miR-7a.2 is more abundant than the CoV2-miR-7a.1 (Fig. S3B-C). Next, we performed RT-qPCR assay to detect the CoV2-miR-7a.1 and CoV2-miR-7a.2 in DICER1-depleted cells. Our results showed that the depletion of DICER1 mRNA (Fig. S3D) was sufficient to reduce the levels of CoV2-miR-7a.1 and CoV2-miR-7a.2 (Fig. 4A), and their reduction was similar to the reduction observed for the canonical dicer-dependent human miR-let-7a (Fig. 4A). These results confirm that CoV2-miR-7a. 1 and CoV2-miR-7a.2 are two isoforms produced by the host enzyme Dicer.

**Figure 4:**
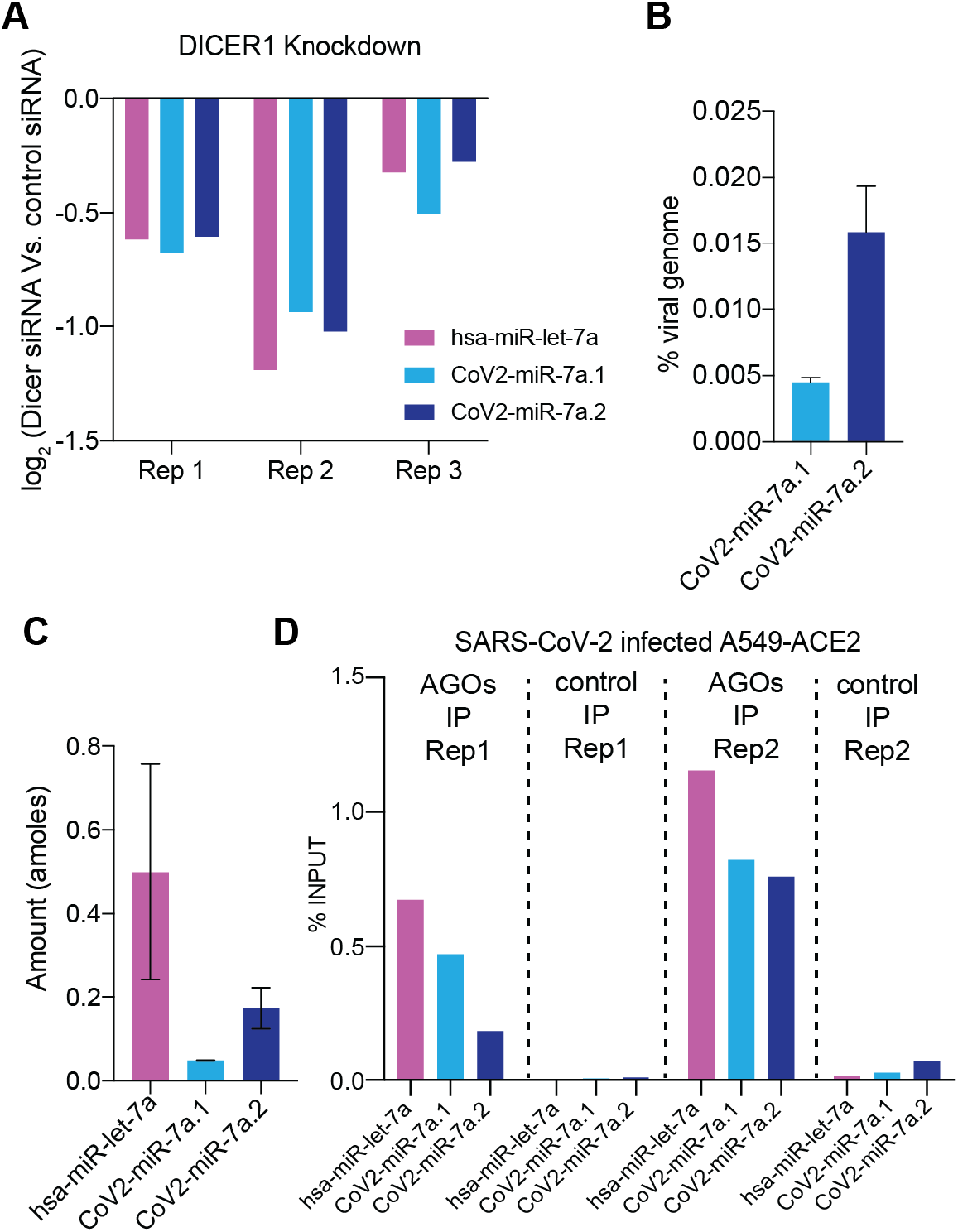
The SARS-CoV-2 miR-7a produces two isoforms processed by DICER and loaded onto AGOs. **(A)** Log_2_ fold changes of the levels of hsa-miR-let-7a, CoV2-miR-7a. 1 and COV2-miR-7a.2 in DICER1 knockdown SARS-CoV-2-infected A549-ACE2 cells compared to control cells analyzed by stem-loop RT-qPCR. Results from three independent replicates are shown. **(B)** Expression levels of the virus-encoded miRNAs as a percentage of the viral genome. Absolute quantification of virus-encoded miRNAs and viral genome from infected A549-ACE2 cells was performed using two spike-in (see methods). Bars and error bars represent the average and standard deviation from two independent experiments. **(C)** Amount of hsa-miR-let-7a, CoV2-miR-7a.1 and CoV2-miR-7a.2 in infected A549-ACE2 cells quantified using a small RNA spike-in of the known amount by stem-loop RT-qPCR. Levels of hsa-miR-let-7a were normalized for the percentage of infection. Bars and error bars represent the average and standard deviation from two independent experiments. **(D)** Loading of hsa-miR-let-7a, CoV2-miR-7a.1 and COV2-miR-7a.2 into AGOs as measured by stem-loop RT-qPCR and analyzed as a percentage of input from the immunoprecipitates (IPs) of either pan-AGO IP or control IgG IP from infected A549-ACE2 cells. Levels of hsa-miR-let-7a were normalized for the percentage of infection.

The processing of miRNAs directly from the viral RNA genome might be a strategy adopted by the host to reduce viral replication and could also explain why virus-encoded miRNAs are not prevalent in RNA-viruses infected cells (Aguado & tenOever, 2018). However, when we quantitatively measured the amount of CoV2-miR-7a produced from the SARS-CoV-2 genome, we found that only 0.01% of the viral genome is used to produce the predominant isoform of CoV2-miR-7a, suggesting that it is unlikely that their production reduces viral genome copy numbers (Fig. 4B). Nonetheless, the amount of the CoV2-miR-7a.2 isoform was comparable with one of the most abundant human miRNAs in infected A549-ACE2 cells, the hsa-miR-let-7a (Fig. 4C, Table S2). Given the abundance of CoV2-miR-7a in infected human cells, we explored the possibility that the CoV2-miR-7a hijacks the human AGOs to regulate host transcripts. To test this hypothesis, we performed RNA immunoprecipitation experiments in A549-ACE2 cells using a pan-AGO antibody that recognizes all four human AGOs (Nelson *et al*, 2007). Our results demonstrated the loading of CoV2-miR-7a.1 and CoV2-miR-7a.2 into human AGOs, with similar efficiency than the human miR-let-7a (Fig. 4D). We confirmed these results by RNA immunoprecipitation experiments performed in infected Caco-2 cells (Fig. S3E). The processing of viral CoV2-miR-7a by human Dicer and the loading onto human AGOs suggest that CoV2-miR-7a might regulate human genes by hijacking the host miRNA machinery.

To investigate whether the CoV2-miR-7a targets human genes, we analyzed the human transcriptome for homology to the sequence of CoV2-miR-7a. 1 and CoV2-miR-7a.2. We found a nearly perfect antisense complementarity to the 3’ untranslated regions (3’UTR) of BATF2 (Fig. 5A), which is a transcription factor that plays a major role in innate immunity during viral infection (Murphy *et al*, 2013; Tussiwand *et al*, 2012). Importantly, BATF2 is an interferon-stimulated gene (ISG) and, as such, its expression is suppressed in numerous human cells infected with SARS-CoV-2 (Blanco-Melo *et al*, 2020), including A549-ACE2 cells. Animal miRNAs target sites that are located in the 3’UTRs of mRNAs and decrease their expression through translation inhibition and mRNA decay (Bartel, 2018). We thus hypothesized that the SARS-CoV-2 miRNA can similarly inhibit the expression of BATF2 mRNA. To test this hypothesis, we transfected A549-ACE2 cells with 22 nt miRNA mimics corresponding to the sequence of CoV2-miR-7a.1, CoV2-miR-7a.2 or a control sequence not targeting the human genome. Given that IFNs induce BATF2 mRNA expression, we performed a timecourse experiment to evaluate the level of expression of ISGs in A549-ACE2 cells at 8, 16, and 24 hours after IFN-α induction compared to non-induced controls (Fig. 6A and Table S3). We found that BATF2 is highly induced by 8 hours of IFN-α treatment and its expression rapidly decays at 16 and 24 hours (Fig. 5B, Fig. 6A). We thus transfected the CoV2-miR-7a.1, CoV2-miR-7a.2 or control mimics in A549-ACE2 cells and analyzed the level of BATF2 mRNAs after 8 hours of IFN-α treatment compared to noninduced control. Our results showed a severe downregulation of BATF2 mRNAs upon 8 hours of IFN-α treatment in cells transfected with CoV2-miR-7a.2, but not in cells transfected with the CoV2-miR-7a. 1 mimic (Fig. 5C). Given that the repressive function of miRNAs is achieved through the base complementarity at their 5’ position, between the 2nd and the 7th nucleotides-the seed region – the imperfect complementarity of CoV2-miR-7a. 1 and BATF2 3’UTR in this region might explain the lack of repression despite the overall extensive complementarity (Fig. 5A, C). These results suggest that CoV2-miR-7a.2, which is the predominant CoV2-miR-7a isoform, specifically interferes with the expression of BATF2 mRNA during SARS-CoV-2 infection through a mechanism similar to the one used by the host miRNA pathway to regulate mRNA targets.

**Figure 5:**
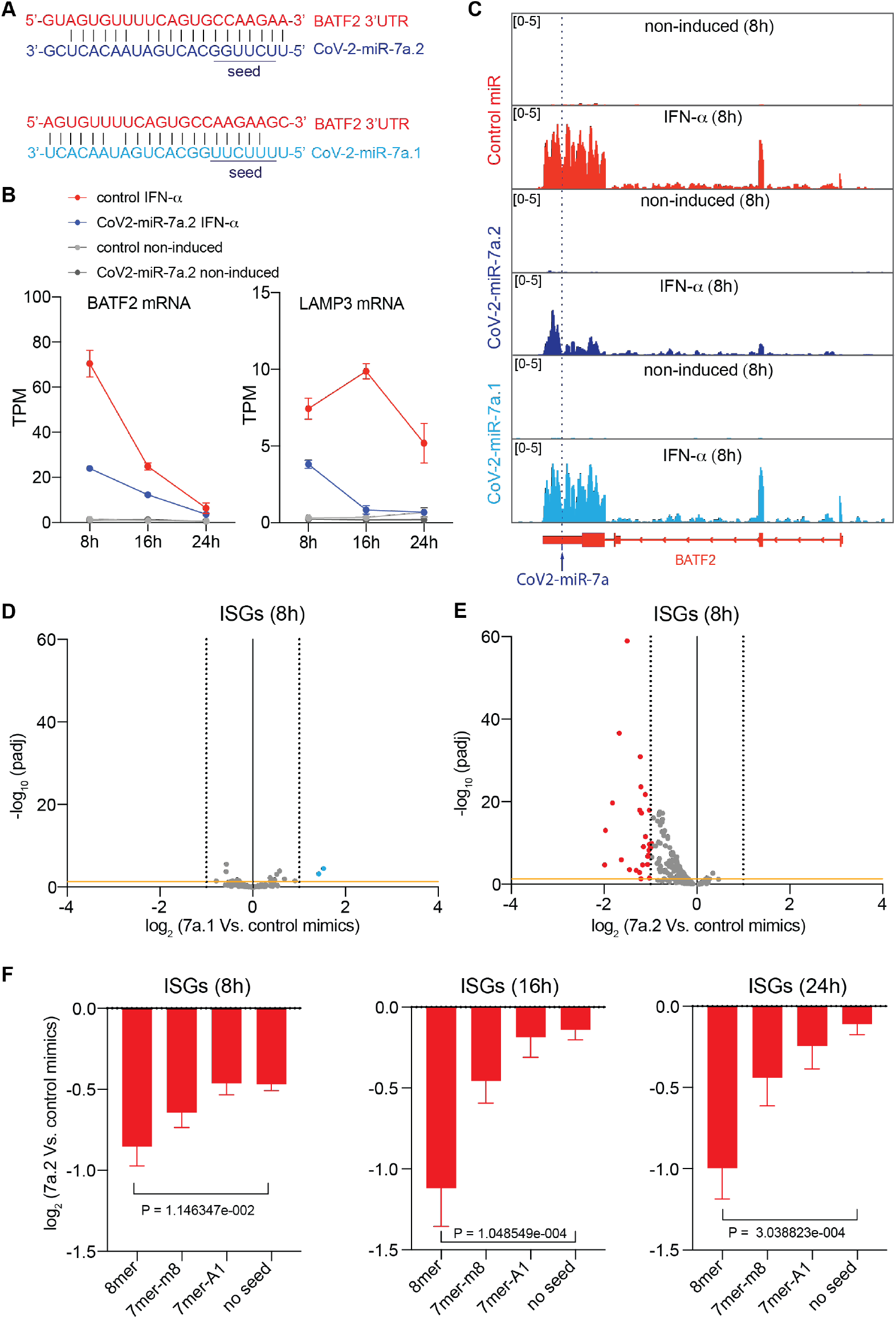
SARS-CoV-2 miR-7a.2 represses the activation of BATF2 and other interferon-stimulated genes. **(A)** Base-pairing of CoV2-miR-7a.2 to complementary 3’UTR site of BATF2 mRNA. The seed region required for binding of miRNAs with its target is underlined. One mismatch in the seed region of CoV2-miR-7a.1 is observed. **(B)** Kinetics of BATF2 and LAMP3 mRNA levels in non-induced and interferon IFN-α-induced (for 8, 16, and 24 h) A549-ACE2 cells transfected with either control or CoV2-miR-7a.2 mimics. Median levels and 95% confident interval of normalized read abundances in transcript per million (TPM) are shown. (n=2). **(C)** Genomic view of the human BATF2 gene showing normalized RNA-seq reads from non-induced and IFN-α-induced (for 8 h) A549-ACE2 cells transfected with either control, CoV2-miR-7a.1 or CoV2-miR-7a.2 mimics. The complementary site of CoV2-miR-7a to BATF2 3’UTR is shown. **(D, E)** Volcano plots showing the log_2_ fold change and corresponding significance levels of ISGs upon 8 hours of IFN-α treatment in A549-ACE2 cells transfected with CoV2-miR-7a. 1 (D) or CoV2-miR-7a.2 (E) compared to control mimic. Significantly downregulated genes are marked in red, and upregulated genes in blue. The orange horizontal line indicates two-tailed P = 0.05. n=2. **(F)** Log_2_ fold change of ISGs as in (E) at 8, 16, 24 hours of IFN-α treatment and categorized based on CoV2-miR-7a.2 target sites 8mer, 7mer-m8, 7mer-A1, and no seed. The mean and standard error of the mean is shown. The two-tailed p values were calculated using the Mann-Whitney-Wilcoxon test. ISGs were calculated as all the upregulated genes (≥ 3-fold; padj < 0.05) in IFN-induced versus non-induced conditions in all time points.

Because miRNAs require only a few nucleotides to be perfectly complementary to the 3’UTR target, the CoV2-miR-7a may potentially regulate other ISGs. We thus analyzed the changes in mRNAs of ISGs at 8 hours of IFN-α treatment compared to non-induced controls in the presence of CoV2-miR-7a.1, CoV2-miR-7a.2 or the control sequence. Notably, we observed a global downregulation of ISGs in the presence of CoV2-miR-7a.2, whereas the CoV2-miR-7a.1 showed negligible effects on the expression of ISGs (Fig. 5D, E). Even though the CoV2-miR-7a.1 does not affect the expression of ISGs in the tested cell lines, we cannot rule out its effect on gene expression in other conditions. Our time-course experiment using IFN-α treatment also showed that some ISGs displayed a more stable expression compared to BATF2, which rapidly decay after induction by IFN-α treatment (Fig. 5B, and Fig. 6A). Therefore, we tested whether the transfection of the CoV2-miR-7a.2 mimic also shows inhibitory effects on ISGs at later time points of the IFN-α treatment. Indeed, our experiment revealed increased global downregulation of ISGs at later time points upon IFN-α stimulation in the presence of CoV2-miR-7a.2 (Fig. 5F and Fig. 6B). Moreover, the level of downregulation correlated with the degree of complementarity of the extended seed region – nucleotides 2-8 and the presence of an adenosine across from the first nucleotide of the miRNAs – (Fig. 5F), similarly to what has been documented for canonical miRNA sites (Bartel, 2018). Indeed, the expression of one of the CoV2-miR-7a.2 ISG targets with the most extended complementarity, the dendritic cell lysosomal associated membrane glycoprotein LAMP3, was almost completely suppressed in the presence of CoV2-miR-7a.2 at later time points (Fig. 5B, Fig. 6B and Fig. S4A). Our sequencing results were confirmed by RT-qPCR analyses on selected CoV2-miR-7a.2 ISG targets at different time points of IFN-α treatment (Fig. S4B, C). Overall, these results suggest that the predominant isoform of CoV2-miR-7a can inhibit the expression of ISGs and thus facilitate SARS-CoV-2 replication.

**Figure 6:**
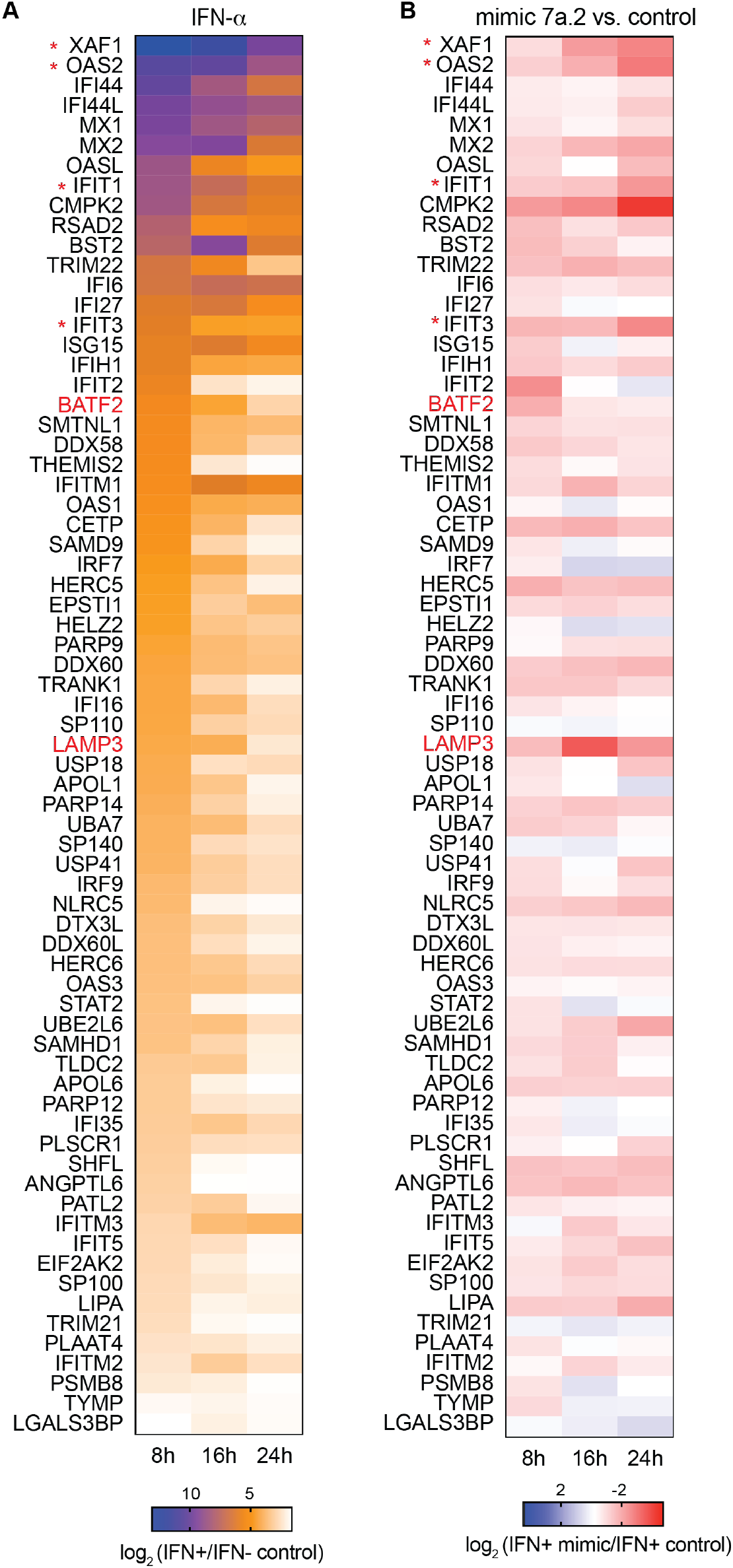
Dynamic expression of ISGs upon type I IFN-a treatment at different time points in A549-ACE-2 cells and in the presence of CoV2-miR-7a.2 mimic. **(A)** Heat map showing log_2_ fold change of expression of ISGs in A549-ACE2 cells across 8, 16 and 24 h time points upon IFN-a treatment compared to non-treated controls. The common upregulated genes (≥ 3-fold; padj < 0.05) in IFN-induced versus non-induced conditions at all time points were categorized as ISGs. **(B)** Heat map showing log_2_ fold change of expression of ISGs across 8, 16 and 24 h time points upon IFN-a treatment in A549-ACE2 cells transfected with CoV2-miR-7a.2 mimic compared to control mimic. Genes marked in red and * indicate the ISGs detected by RT-qPCR in Fig. S4.

The detection of the CoV2-miR-7a.1 and CoV2-miR-7a.2 during infection of human cell lines does not provide evidence for their processing in more physiological conditions. We first evaluated the processing of the CoV2-miR-7a.1 and CoV2-miR-7a.2 using 2D human colon organoids, which are a relevant model to study SARS-CoV-2 biology and infection (Stanifer *et al*, 2020). Using this system, we were able to detect by RT-qPCR both CoV2-miR-7a.1 and CoV2-miR-7a.2 during SARS-CoV-2 infection at low and high multiplicity of infection (MOI) (Fig. S5A, B). Moreover, as we observed in cell lines (Fig. S3B-C), the CoV2-miR-7a.2 was the predominant isoform of the CoV2-miR-7a in infected human intestinal 2Dorganoids (Fig. S5A, B).

To ensure that the observed processing of CoV2-miR-7a does not result from a byproduct of *in vitro* cell culture, we tested the presence of CoV2-miR-7a.1 and CoV2-miR-7a.2 in COVID-19 patients. We extracted small RNAs from nasopharyngeal swab samples collected from patients who tested positive for the presence of SARS-CoV-2 or from patients infected with seasonal human coronaviruses (HCoV). RT-qPCR assays revealed the presence of the two CoV2-miR-7a. 1 and CoV2-miR-7a.2 isoforms exclusively in COVID-19 patients but not in HCoV-infected patients (Fig. 7A), while the human miR-let-7a was readily detected in all patients (Fig. S5C). Moreover, we failed to detect small RNA from the proximal ORF6 region, indicating that the amplification of the two isoforms of CoV2-miR-7a is not the result of genomic viral RNA degradation in nasopharyngeal swab samples (Fig. 7A). In addition, the relative expression of the CoV2-miR-7a.1 and CoV2-miR-7a.2 among different COVID-19 patients correlated with genomic viral RNA levels detected in the swab samples (Fig. 7A), suggesting that the higher is the abundance of the SARS-CoV-2 genome, the higher is the processing of CoV2-miR-7a.1 and CoV2-miR-7a.2 by the human Dicer during viral replication in the upper respiratory tract. The relative levels of CoV2-miR-7a.2 compared to CoV2-miR-7a.1 confirmed that the CoV2-miR-7a.2 is the predominant isoform of CoV2-miR-7a in COVID-19 patients (Fig. 7A). Finally, we selected three nasopharyngeal swab samples collected from patients with high viral load to perform small RNA sequencing. Even though the majority of reads were RNA degradation products, we were able to identify 22nt small RNA reads mapping to the CoV2-miR-7a genomic regions (Fig. 7B). This result confirms the production of CoV2-miR-7a in human patients infected with SARS-CoV-2.

**Figure 7:**
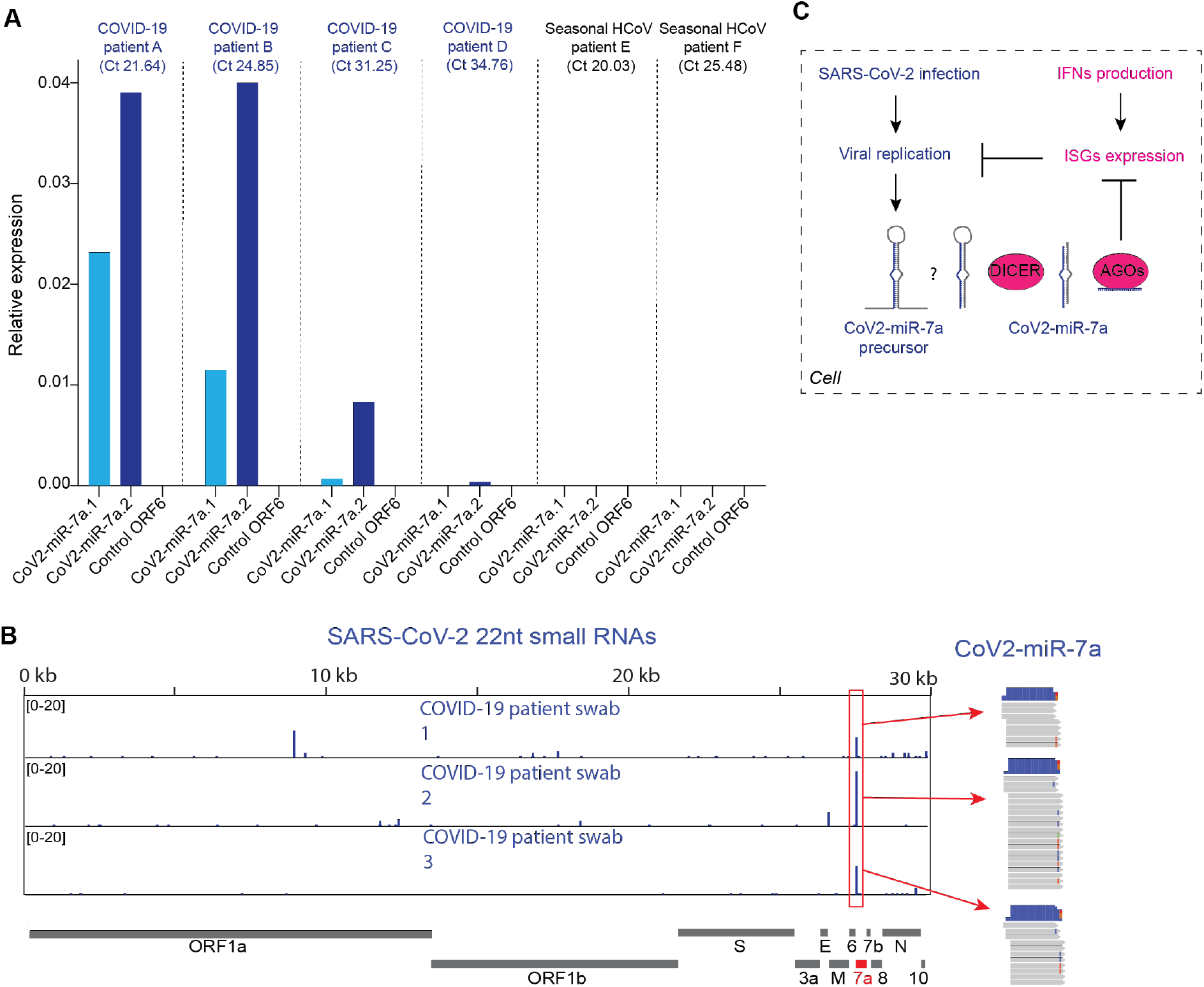
The two CoV2-miR-7a isoforms are produced in COVID-19 patients. **(A)** Expression levels of CoV2-miR-7a.1, CoV2-miR-7a.2 and a 22 nt region from the ORF6 of the viral genome that does not produce detectable levels of small RNAs (Control ORF6) analyzed by stem-loop RT-qPCR from nasopharyngeal swabs of patients tested positive for COVID-19 or another seasonal human coronavirus (HCoV). Relative expression to hsa-miR-let-7a is shown. Ct values in parenthesis refer to the Ct value for the detection of viral genome in patient swab samples. **(B)** SARS-CoV-2 genomic view showing the distribution of normalized 22nt small RNA reads from nasopharyngeal swabs of three patients tested positive for COVID-19. The most abundant small RNAs are marked by the red boxes and correspond to the CoV2-miR-7a. **(C)** Model for the production and function of Cov2-miR-7a in COVID-19 patients.

## DISCUSSION

Our study identified the first *bona fide* miRNA encoded by the SARS-CoV-2 genome, which is processed by the host miRNA pathway and loaded by human AGOs. It also represents the first evidence of a genuine miRNA encoded by a cytoplasmic RNA virus. Our data suggests that SARS-CoV-2 uses a viral miRNA to hijack the host miRNA machinery and act like host miRNAs to repress the expression of ISGs through sequence complementarity to sites located in their 3’UTR. We propose that viral miRNA production is one of the mechanisms used by the virus to repress ISGs and evade the innate immune response (Fig. 7C). Therefore, CoV2-miR-7a might contribute to the documented impaired activation of ISGs upon SARS-CoV-2 infection (Blanco-Melo *et al*, 2020; Kim & Shin, 2021).

One of the hallmarks of severe COVID-19 patients is the decreased expression of ISGs accompanied by low levels of Type I IFN levels and high blood viral load (Hadjadj *et al*, 2020). Among the ISGs targeted by the CoV2-miR-7a.2, two of the most regulated targets, BATF2, which plays a fundamental role during viral infections (Murphy *et al*, 2013), and LAMP3, which inhibits influenza virus replication (Zhou *et al*, 2011), are both required for dendritic cell function in adaptive immunity (Tussiwand *et al*, 2012; Saint-Vis *et al*, 1998). Given that acute SARS-CoV-2 infection impairs dendritic cell response (Zhou *et al*, 2020c), we speculate that the suppression of key ISGs, including BATF2 and LAMP3, by the CoV2-miR-7a might constitute one of the mechanisms responsible for the reduced dendritic cell response in patients with severe COVID-19 disease. The sequence of genomic region encoding CoV2-miR-7a has been evolutionarily conserved in different variants of the SARS-CoV-2 and is less prone to mutations than adjoining genomic region, highlighting its biological relevance for viral infection.

One limitation of this study is that we could not evaluate whether the presence and abundance of the CoV2-miR-7a.2 correlates with disease progression or with impaired ISGs activation in patients with severe COVID-19 disease outcome. Thus, future studies will address the relevance of the CoV2-miR-7a in the progression of the COVID-19 disease. Furthermore, the recent evolution of the CoV2-miR-7a sequence in the SARS-CoV-2 genome may facilitate the development of specific therapeutic approaches to potentially target and dampen the virulence of SARS-CoV-2 infection in COVID-19 patients.

## MATERIALS AND METHODS

### Ethical statement

Samples used in this study were collected as part of approved ongoing surveillance conducted by the National Reference Center for Respiratory Viruses (NRC) at Institut Pasteur (WHO reference laboratory providing confirmatory testing for COVID-19). The investigations were carried out in accordance with the General Data Protection Regulation (Regulation (EU) 2016/679 and Directive 95/46/EC) and the French data protection law (Law 78-17 on 06/01/1978 and Décret 2019-536 on 29/05/2019).

### Human swab sample collection

For each suspected COVID-19 case, respiratory samples from the upper respiratory tract (nasopharyngeal swabs) were sent to the NRC to perform SARS-CoV-2-specific real-time RT-PCR.

### Cell culture

Human lung A549-ACE2 cells, which have been modified to stably express ACE2 via lentiviral transduction, were generated in the laboratory of Pr. Olivier Schwartz, (Institut Pasteur, Paris, France). Human colorectal adenocarcinoma Caco-2 and African green monkey Vero E6 cells were purchased from ATCC. A549-ACE2, Caco-2 and Vero E6 cells were cultured in high-glucose DMEM media (Gibco) supplemented with 10% Fetal Bovine Serum (FBS; Sigma) and 1% penicillin-streptomycin (P/S; Gibco). Cells were maintained at 37°C in a humidified atmosphere with 5% CO_2_.

### Virus and infections

The strain BetaCoV/France/IDF0372/2020 was supplied by the National Reference Centre for Respiratory Viruses hosted by Institut Pasteur (Paris, France) and headed by Pr. S. van der Werf. The human sample from which the strain was isolated has been provided by Dr. X. Lescure and Pr. Y. Yazdanpanah from the Bichat Hospital, Paris, France. Viral stocks were produced by amplification on Vero E6 cells, for 72 h in DMEM 2% FBS. The cleared supernatant was stored at −80°C and titrated on Vero E6 cells by using standard plaque assays to measure plaque-forming units per mL (PFU/mL). A549-ACE2 and Caco-2 cells were infected at MOI of 3 and 0,3, respectively, in DMEM without FBS. After 2 h, DMEM with 5% FBS was added to the cells. 48 h post-infection, cells were lysed using TRIzol LS reagent (Thermo Fischer Scientific) and RNA was extracted following manufacturer’s instructions or cells were lysed using FA buffer (50 mM HEPES pH 7.5, 1 mM EDTA, 1 % Triton X-100, 0.1 % sodium deoxycholate, 150 mM NaCl, RNase inhibitor 40 U/mL, Halt™ Protease inhibitor cocktail 1x) for immunoprecipitation experiments. For measurement of miRNAs in the supernatant, RNA was isolated from the supernatant of infected cells using TRIzol LS reagent.

### Analysis of infected cells by flow cytometry

Flow cytometry analyses were performed for each experiment to evaluate the percentage of infected cells. Cells were fixed in 4% PFA for 20 minutes at 4°C and intracellular staining was performed in PBS, 1% BSA, 0.05% sodium azide, 0.05% Saponin. Cells were incubated with antibodies recognizing the spike protein of SARS-CoV-2 (anti-S2 H2 162, a kind gift from Dr. Hugo Mouquet, Institut Pasteur, Paris, France) for 30 minutes at 4°C, and then with secondary antibodies (anti-human-Alexa Fluor-647) for 30 minutes at 4°C. Cells were acquired on an Attune NxT Flow Cytometer (Thermo Fisher) and data analyzed with FlowJo software.

### Transfection of miRNA mimics

1 nM of CoV2-miR-7a.1, CoV2-miR-7a.2, or control mimics (Invitrogen) were transfected in A549-ACE2 or Caco-2 cells using Lipofectamine RNAiMax (Thermo Fischer Scientific). 24 h post-transfection, cells were treated with or without 100U of human Interferon Alpha 2 (PBL Assay Science) for 8, 16 or 24 hours. Cells were then lysed using TRIzol LS reagent (Thermo Fischer Scientific) and RNA was extracted following the manufacturer’s instructions or lysed in RIPA buffer (Thermo Fischer Scientific) for western blot analysis.

### siRNA-mediated knockdown

A549-ACE2 cells were transfected using Lipofectamine RNAiMax (Life Technologies) with 10 nM of control (#4390843, Ambion) or DICER1 siRNAs (#4390824, Ambion), following the manufacturer’s instructions. 48 h after transfection, cells were infected with SARS-CoV-2 for 24 h and then lysed using TRIzol LS reagent (Thermo Fischer Scientific).

### Infection of 2D colon organoids with SARS-CoV-2

Human tissues were a kind gift of by Pr. Iradj Sobhani (Département de Gastroentérologie, Hôpital Henri Mondor, Créteil). They were collected from surgical resection in accordance with the recommendations of the Hospital. Human colonic organoid cultures were generated from isolated crypts or from frozen tissues. They were maintained in culture for expansion prior to dissociation and plating on 0.4 μm pore polyester membrane of Transwell^®^ inserts (Corning). Organoids were then recovered from the Matrigel using Cell recovery solution (Corning) and multiple steps of pipetting. After centrifugation, the organoid fragments were washed in cold DMEM and re-suspended in the organoid medium. 2Dorganoids were seeded in chambers pre-coated with 50 μg/ml human collagen IV (Millipore) for 2 h at 37°C. 700 μl of the medium was added to the bottom well of the chamber and the cells were incubated at 37°, 5% CO_2_ for 2 days before changing the medium to the top compartment. The confluency of the monolayer was reached after 3-4 days of culture and differentiation was induced for 4 days. Infection was performed as described above, by washing the cells after 2 h of virus incubation.

### Immunoprecipitation

A549-ACE2 and Caco-2 cells, infected and not, were lysed in FA buffer as described above. From the total lysate (~0.5 mg/ml), 10 % lysate was saved as input and IP was performed using an anti-pan-AGO antibody (clone 2A8, MABE56 Sigma-Aldrich) and an anti-FLAG M2 antibody (F3165, Sigma-Aldrich) was used for control IPs. The antibody was incubated with the lysates overnight at 4°C, followed by antibody capture by 40 μl Dynabeads™ Protein G (10003D, Invitrogen) for 3 h at 4°C on a rotor. Beads were then captured on a magnetic stand and washed four times with the FA buffer for 10 min at 4°C. After the final wash, the beads were captured on a magnetic stand and total RNA from input and the immunoprecipitate bound on beads was extracted using TRIzol™ Reagent as per manufacturer’s instruction.

### RNA extraction

For total RNA extraction, infected or non-infected cells were directly lysed with TRIzol™ LS (Invitrogen,) and total RNA was isolated according to the manufacturer’s instructions. For RNA-seq or RT-qPCR analysis, a maximum of 10 μg total RNAs was treated with 2 U Turbo DNase (Ambion) at 37 °C for 30 min followed by acid phenol extraction and ethanol precipitation. An Agilent 2200 TapeStation System was used to evaluate the RIN indexes of all RNA preps, and only samples with RNA integrity number (RIN) > 8 were used for further investigations.

### RT-qPCR

Reverse transcription for total RNA was performed using random hexamer primers according to the manufacturer’s instructions using M-MLV reverse transcriptase (Invitrogen, Ref. 28025013). For RT of sRNA’s, specific RT primers were used (Table S4). Quantitative PCR (qPCR) was carried out using Applied Biosystems Power up SYBR Green PCR Master mix following the manufacturer’s instructions and using an Applied Biosystems QuantStudio 3 Real-Time PCR System. Primers used for qPCR are listed in Table S4.

For absolute quantification of viral miRNAs in comparison for viral genome 1:1000 dilution of ERCC RNA Spike-In Mix (ERCC-130 12 amoles) (Invitrogen, 4456740) and a custom sRNA oligo (GAGAGCAGUGGCUGGUUGAGAUUUAAU, 8 nmoles) were added to total RNA prior to RT. Known amounts of the spike-ins were used for quantification of viral genome and miRNAs. Levels of hsa-miR-let-7a were normalised to viral miRNAs as a ratio of total hsa-miR-let-7a to the infection rate of infected cells.

For immunoprecipitation experiments, levels of miRNAs were expressed as a percentage of input. For all other fold change comparisons, details are provided in figure legends.

### Small RNA-seq

Total RNA (2-5 μg) was resolved on a 15 % TBE-Urea gel (Invitrogen EC6885BOX). RNA of size between 17-25 nt was excised from the gel and extracted in 0.3 M NaCl overnight at 25°C. Size selected RNA was used to prepare libraries following previously described methodology (Barucci *et al*, 2020), which included the ligation of 3’end and 5’end adaptors each having four randomized nucleotides to minimize ligation biases. The randomized 8nt were also used to remove possible PCR duplicates occurring in the PCR amplification step of the library preparation. We also exclusively ligated monophosphate small RNAs with a pre-adenylated 3’ adaptor. Libraries were multiplexed and their quality was assessed on TapeStation (Agilent). They were quantified using the Qubit Fluorometer High Sensitivity dsDNA assay kit (Thermo Fisher Scientific, Q32851) and sequenced on a NextSeq-500 Illumina platform using the NextSeq 500/550 High Output v2 kit 75 cycles (FC-404-2005).

### Strand-specific RNA-seq library preparation

DNAse-treated RNA with high RIN value was used to deplete ribosomal RNA using NEBNext^®^ rRNA Depletion Kit (Human/Mouse/Rat) (NEB #E6350) as per manufacturer’s instructions. Strand-specific RNA libraries were prepared using at least 100 ng of rRNA depleted RNAs using NEBNext Ultra II Directional RNA Library Prep Kit for Illumina (E7760S) as per the manufacturer’s instructions.

### Data Analysis

#### RNA-seq

Multiplexed Illumina sequencing data were demultiplexed using Illumina bcl2fastq converter (version 2.20.0.422). Reads were aligned on the Homo sapiens genome (Build version GRCh38, NCBI) using Hisat2 (Kim *et al*, 2015) (version 2.2.1) with the default settings. After alignment, reads mapping to annotated protein-coding genes were counted using featureCounts (version 2.0.1). Annotations were obtained from the Ensembl release 100. Counted reads for protein-coding genes were used for differential expression analysis using the R/Bioconductor package DESeq2 (Love *et al*, 2014) (version 1.26.0).

#### Small RNA-seq

Multiplexed Illumina sequencing data were demultiplexed using Illumina bcl2fastq converter (version 2.20.0.422). The 3’ adapter was trimmed from raw reads using Cutadapt (Martin, 2011) v.1.15 using the following parameter: -a TGGAATTCTCGGGTGCCAAGG --discard-untrimmed. 5’ and 3’ end unique molecular identifiers (UMIs) were used to deduplicate the trimmed reads. Deduplication was performed by first sorting reads by sequence using the option -s in fastq-sort (from fastq-tools v.0.8; https://github.com/dcjones/fastq-tools/tree/v0.8) and then using a custom Haskell program that retained the best quality reads at each position among reads of identical sequences. Then, 4-nucleotide UMIs were trimmed at both ends using Cutadapt (options, -u 4 and -u -4). Finally, we selected only deduplicated reads ranging from 18 to 26 nucleotides using bioawk (https://github.com/lh3/bioawk). The selected 18-26-nucleotide reads were aligned on the SARS-CoV-2 genome (assembly Jan.2020/NC_045512v2, UCSC) or on the Homo sapiens genome (Build version GRCh38, NCBI) using Bowtie2 (Langmead & Salzberg, 2012; Li *et al*, 2009) v.2.4.2 with the following parameters: -L 6 -i S,1,0.8 -N 0. Mapped reads were divided in sense and antisense reads in respect to the reference genome using samtools (Li *et al*, 2009) (version 1.3.1) while 22-nucleotide reads were extracted from mapped reads using bioawk. The size distribution of all categories of mapped reads was calculated using bioawk. Reads mapping to CoV-2-microRNAs were counted using a custom script. First, a bed file with the coordinates of all the putative 22-nucleotide RNAs encoded by the SARS-CoV-2 genome was created and used to extract their sequences using bedtools. Second, the occurrence of each putative 22-nucleotide RNA among aligned reads was counted.

#### Generation of bigwig files

For RNA-seq libraries, normalized bigwig files were generated from the mapping results using CPM as a normalization factor. This normalized coverage information was computed for 20 bp bins using the bamCoverage from deeptools (version 3.5.0).

For small RNA-seq libraries, normalized bigwig files were generated from the mapping results using the sum of the total number of reads mapping on the SARS-Cov-2 genome or Homo sapiens genome as a normalization factor. This normalized coverage information was computed for 20 bp bins using the bamCoverage from deeptools (version 3.5.0).

#### Size distribution

Size distribution of mapped reads was computed using bioawk. To compute the size distribution around a specific region, reads mapping to a specific region of the genome were extracted using samtools and size distribution was computed using bioawk.

#### Identification of CoV2-miR-7a.2 complementary sites on human 3’UTR

The sequences of the 3’UTR of human genes have been retrieved using the Ensembl BioMart (database Ensembl Genes 101 – Human genes (GRCh38.p13)). Genes were divided into three different categories based on the presence in their 3’UTR of the miRNA complementary sites 8mer, 7mer-m8 or 7mer-A1 as described in (Bartel, 2018) (Table S5).

#### RNA folding structure

RNA secondary structure of the CoV2-miR-7a precursor region has been determined with the Vienna RNA Package (Hofacker, 2003) using the first 70 nt of the open reading frame of the ORF7a of different SARS coronaviruses.

#### Conservation analysis of viral genomic region encoding CoV2-miR7a

A conservation study was performed on data obtained from GISAID (https://www.epicov.org/). A total of 1,612,294 sequences of SARS-CoV2 obtained between January 2020 and April 2021 were aligned, and the region of ORF7a (from position 27,394 to 27,759) was extracted for all sequences.

All sequences were compared to the original SARS-CoV2 sequence (EPI_ISL_402124) and mutations were counted for all positions of the region. Conservation percentage was calculated as number of correct bases/Total number of sequences. The same calculations were made on subsets of sequences corresponding to the major known variants. Main variants considered includes with sequence counts mentioned in parenthesis for each: B.1.1.214 (16,417), B.1.1.7 (646,208), B.1.1 (41,042), B.1.160 (21,077), B.1.177 (66,212), B.1.2 (71,986), B.1.351 (15,910), B.1.429 (25,413), B.1.526 (16,530), B.1 (63,998), P.1 (18,193).

### Gene lists

Gene lists used in this study are shown in Table S5, which also includes the log_2_ fold changes and padj used for all the analysis presented.

## ACKNOWLEDGMENTS

We thank the members of the Pasteur COVID-19 Task Force, Christophe D’Enfert, Felix Rey and Olivier Schwartz for scientific discussions and support during the realization of this project and Geneviève Millon for critical reading of the manuscript. We also thank the Biomics Platform, C2RT, Institut Pasteur, Paris, France, supported by France Génomique (ANR-10-INBS-09-09), IBISA and the Illumina COVID-19 Projects’ offer.

## AUTHOR CONTRIBUTIONS

G.C. identified and developed the core questions addressed in the project, including the identification of the viral miRNA, and wrote the paper with the contribution of all the authors. M.S. performed all the biochemical, molecular biology and sequencing experiments. P.Q. performed all the bioinformatic analysis. M.C. performed most of the cell culture and infection experiments. N.J. supervised the cell culture and infection experiments. L.B. performed the RT-qPCR experiments on human swab samples and on some human cell lines. C.M. performed the conservation analysis of the ORF7a among the SARS-CoV2 sequenced genomes. T.V. and M.V. provided the first batch of infected cell lines. S.v.d.W., S.B, F.D., provided the human swab samples and performed RNA extraction. N.S and G.N performed experiments on infected human 2D-organoids. M.B. helped with some experiments. G.C., M.S., M.C., and N. J., analyzed the results.

## DATA AND MATERIALS AVAILABILITY

All the sequencing data are available at the following accession numbers GSE162318. All other data supporting the findings of this study are available from the corresponding author on reasonable request.

## SUPPLEMENTARY FIGURES

**Figure S1:**
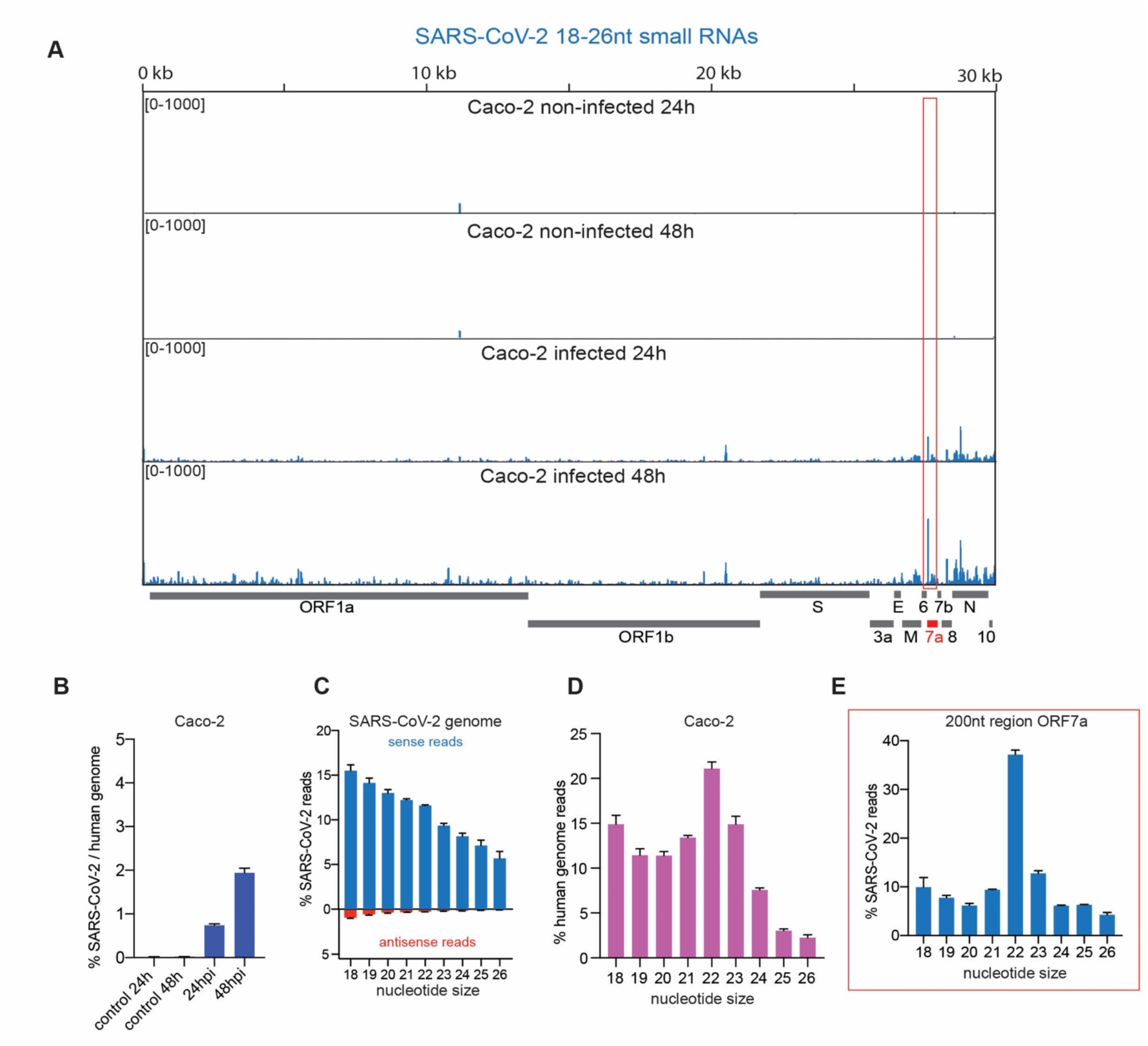
Identification of SARS-CoV-2 miR-7a in A549-ACE2 human cells. **(A)** SARS-CoV-2 genomic view showing the distribution of normalized total small RNA reads (18-26 nt in length) from Caco-2 cells at 24 and 48 hpi and non-infected controls. The red box marks a distinct peak observed in ORF7a that has been further characterized. n=2. **(B)** Percentage of total small RNA reads (18-26 nt) mapping on SARS-Cov-2 genome compared to the human genome from SARS-CoV-2 in Caco-2 cells at 24 and 48 hpi and non-infected controls. The mean and standard deviation of 2 experiments is shown. **(C)** Percentage of the size distribution of SARS-CoV-2 total small RNA sense (blue) and antisense (red) reads from Caco-2 cells at 48 hpi. The mean and standard deviation of the 2 experiments are shown. **(D)** The size distribution of total small RNA reads mapping on the human genome from Caco-2 cells at 48 hpi shows a bias for 22 nt. The mean and standard deviation of 2 experiments are shown. **(E)** Percentage of the size distribution of small RNA reads from Caco-2 cells at 48 hpi that map to the 200 nt region surrounding the distinct small RNA peak identified in ORF7a (red box in panel A). A bias for 22 nt typical of Dicer processed small RNAs is revealed. The mean and standard deviation of 2 experiments are shown.

**Figure S2:**
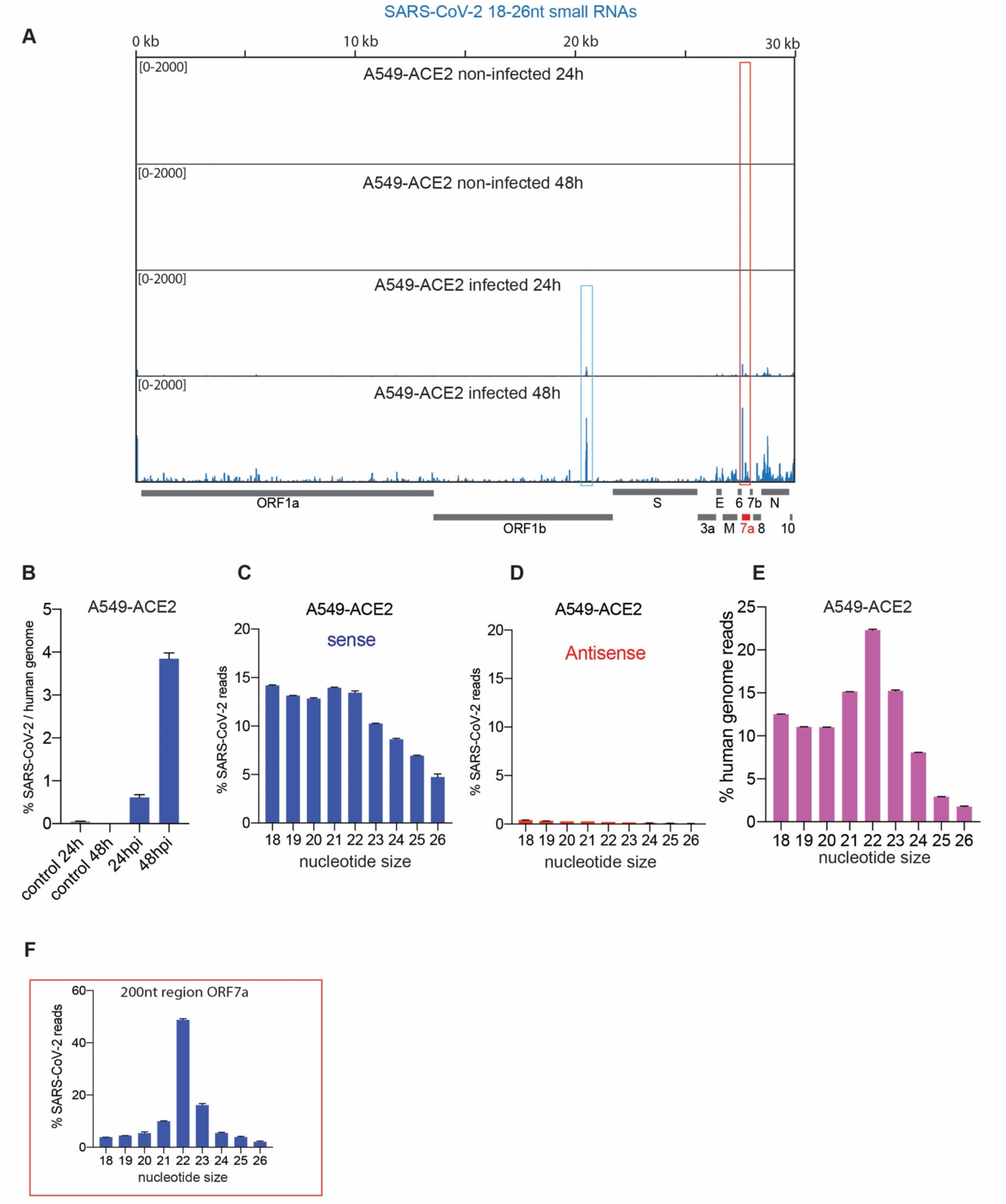
Identification of SARS-CoV-2 miR-7a in A549-ACE2 human cells. **(A)** SARS-CoV-2 genomic view showing the distribution of normalized total small RNA reads from infected A549-ACE2 cells at 24hpi and 48 hpi and non-infected controls. The red box marks a distinct peak observed in ORF7a that has been further characterized. The blue box marks another peak derived from ORF1b, which is not abundant in Caco-2 infected cells. **(B)** Percentage of total small RNA reads (18-26 nt) mapping on SARS-Cov-2 genome compared to the human genome from SARS-CoV-2 in A549-ACE2 cells at 24 and 48 hpi and non-infected controls. The mean and standard deviation of 2 experiments are shown. **(E)** The size distribution of total small RNA reads mapping on the human genome from A549-ACE2 cells at 48 hpi shows a bias for 22 nt. The mean and standard deviation of 2 experiments are shown. **(C-D)** The size distribution of SARS-CoV-2 total small RNA sense (blue) (**C**) and antisense (red) (**D**) reads from A549-ACE2 cells at 48 hpi. The mean and standard deviation of 2 experiments are shown. **(F)** The size distribution of small RNA reads from A549-ACE2 cells at 48 hpi that map to the 200 nt region surrounding the distinct small RNA peak identified in ORF7a (red box in panel E). A bias for 22 nt typical of Dicer processed small RNAs is revealed. The mean and standard deviation of 2 experiments are shown.

**Figure S3:**
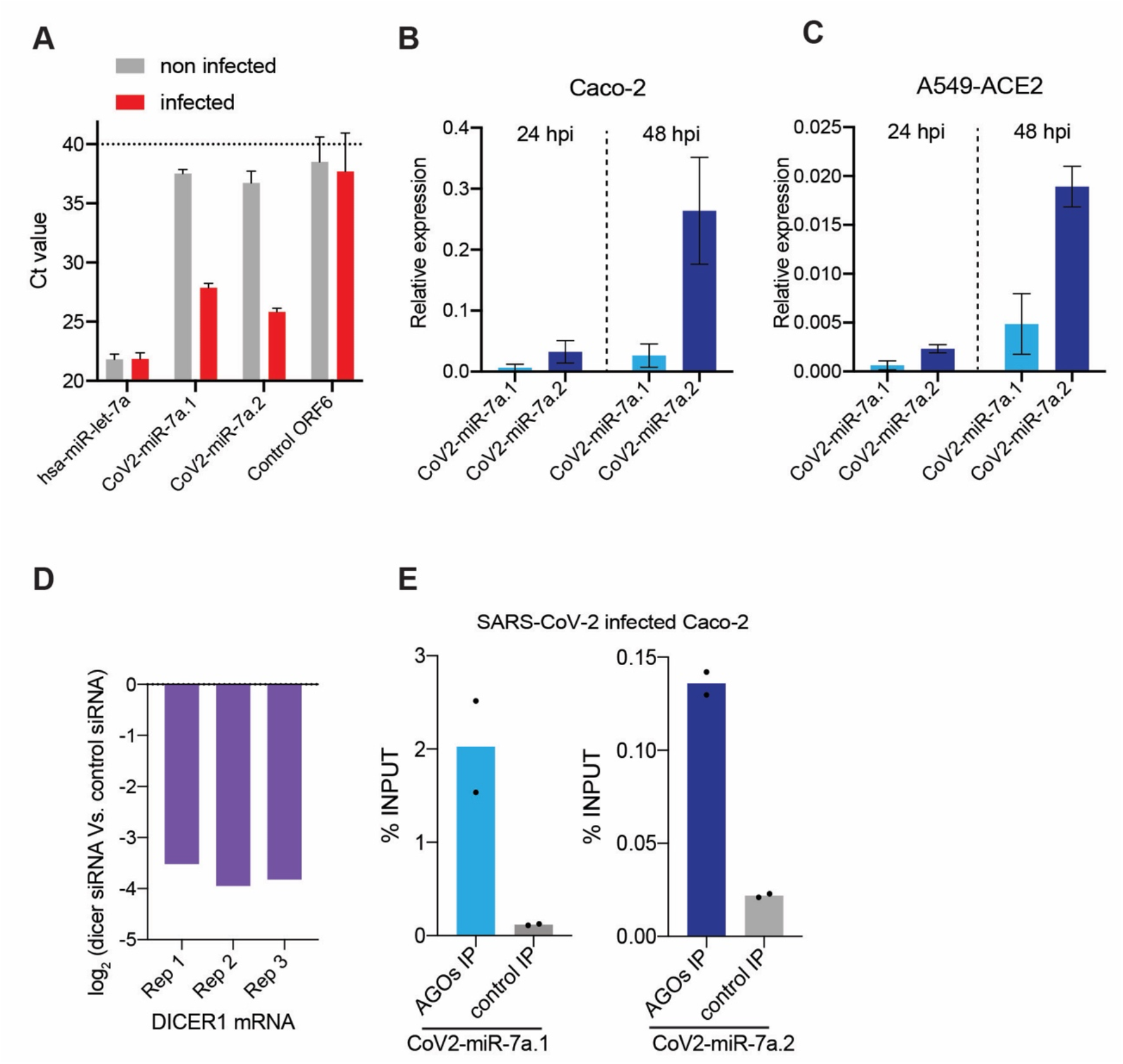
RT-qPCR quantification of SARS-CoV-2 miR-7a in human and loading by AGOs. **(A)** Ct values for hsa-miR-let-7a, CoV2-miR-7a. 1, CoV2-miR-7a.2 and a 22 nt region from the ORF6 of the viral genome that produces a low level of small RNAs (Control ORF6) were determined by stem-loop RT-qPCR performed in A549-ACE2 cells. The mean and standard deviation of 2 experiments are shown. **(B, C)** Expression levels of CoV2-miR-7a. 1 and CoV2-miR-7a.2 by stem-loop RT-qPCR in Caco-2 (B) and A549-ACE2 cells (C) at 24 and 48 hpi. The mean and standard deviation of 2 experiments are shown. Relative expression to hsa-miR-let-7a is shown. **(D)** Levels of DICER1 mRNA were analyzed by RT-qPCR upon siRNA-mediated DICER1 knockdown in A549-ACE2 cells at 48 hpi compared to control siRNAs in the three biological replicates. Actin mRNA was used as internal control **(E)** Loading of CoV2-miR-7a.1 and COV2-miR-7a.2 into AGOs as measured by stem-loop RT-qPCR and analyzed as a percentage of input from the immunoprecipitates (IPs) of either pan-AGO IP or control IgG IP from Caco-2 cells at 48 hpi. n=2.

**Figure S4:**
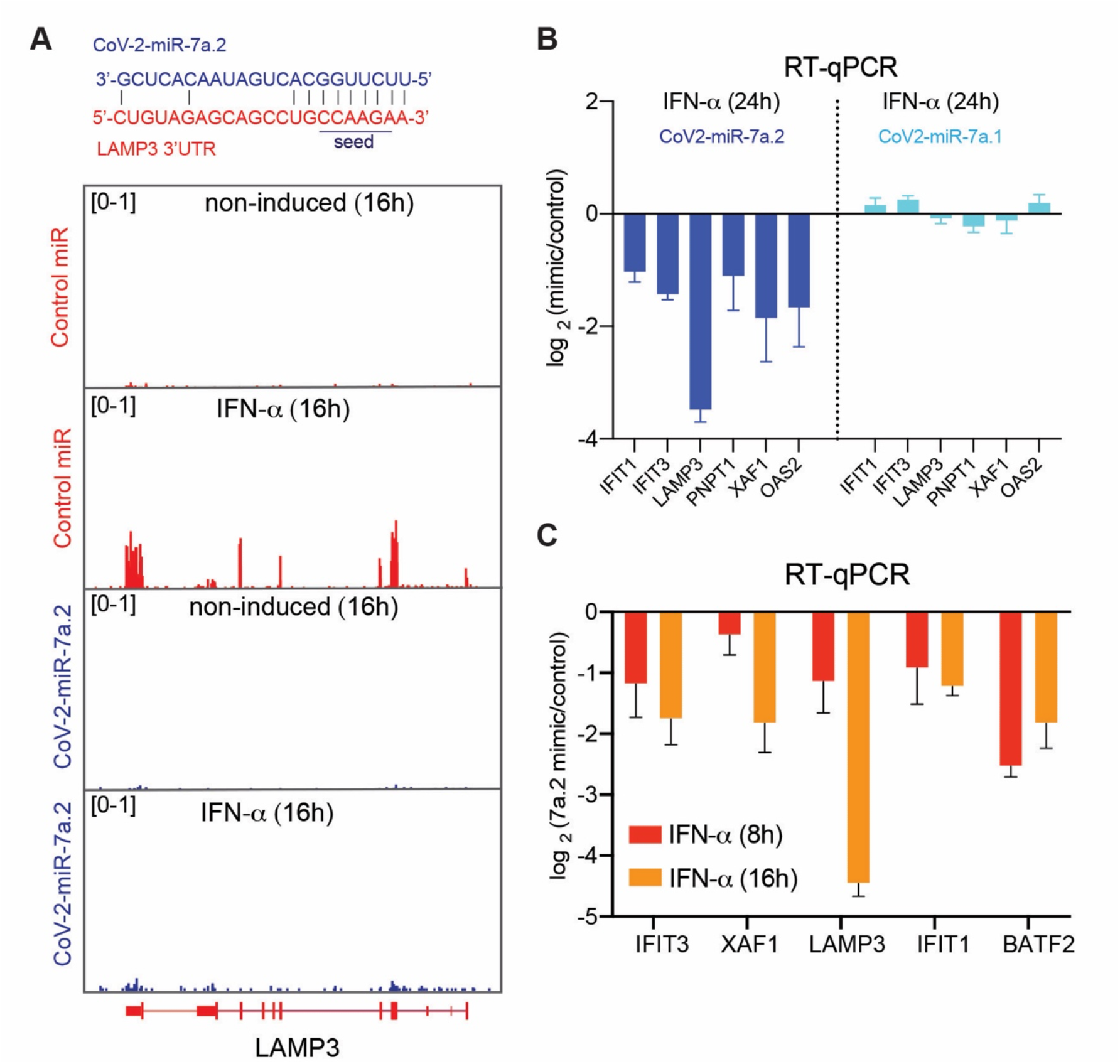
Regulation of ISGs by SARS-CoV-2 miR-7a across different time points of IFN-a treatment. **(A)** Genomic view of the human LAMP3 gene showing normalized RNA-seq reads from non-induced and IFN-α-induced (for 16 h) A549-ACE2 cells transfected with CoV2-miR-7a.2 or control mimics. The base-pairing of CoV2-miR-7a.2 to complementary 3’UTR site of LAMP3 is shown above and the seed region required for binding of miRNAs with the target is underlined. **(B)** log_2_ fold change of expression of selected ISGs measured by RT-qPCR in IFN-a-treated A549-ACE2 cells transfected with CoV2-miR-7a.2 or COV2-miR-7a.1 compared to control mimic at 24 h upon IFN-a treatment. The mean and standard deviation of 3 experiments are shown. **(C)** log_2_ fold change of expression of selected ISGs measured by RT-qPCR in IFN-a-treated A549-ACE2 cells transfected with CoV2-miR-7a.2 mimic compared to control mimic at 8 and 16 h upon IFN-a treatment. The mean and standard deviation of 3 experiments are shown.

**Figure S5:**
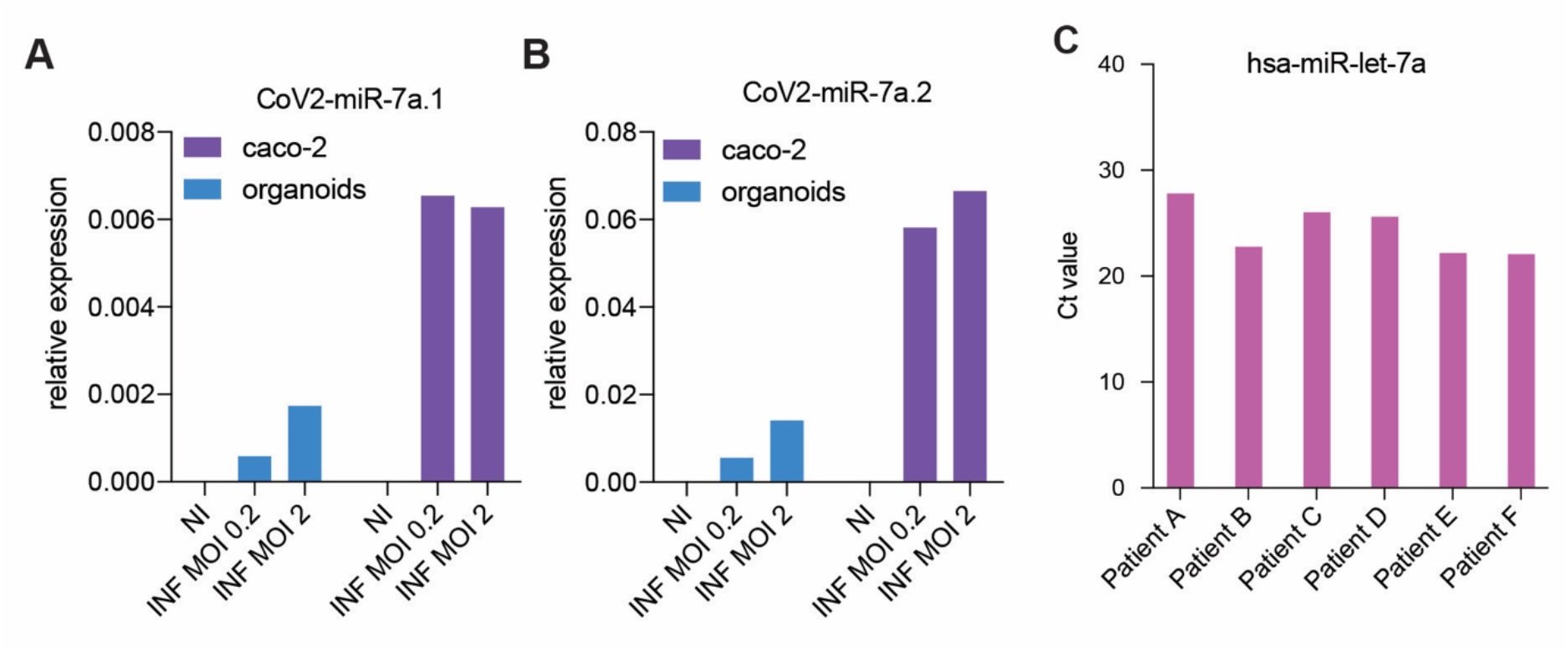
The SARS-CoV-2 miR-7a is detected in 2D human colon organoids and secreted from A549-ACE2 cells. **(A-B)** Detection of CoV2-miR-7a. 1 (A) and CoV2-miR-7a.2 (B) by stem-loop RT-qPCR in no-infected (NI) and SARS-CoV-2-infected (INF) 2D human colon organoids and Caco-2 cells at an MOI of 0.2 and 2. Relative expression to hsa-miR-let-7a is shown. **(C)** Ct values for hsa-miR-let-7a measured by stem-loop RT-qPCR from nasopharyngeal swabs of patients tested positive for COVID-19 or another seasonal HCoV (as in Fig. 7A).

## SUPPLEMENTARY TABLES

**Table S1.** Total small RNA reads mapped on human and SARS-CoV-2 genome.

**Table S2.** Percentage and RPM of human miRNAs.

**Table S3.** log_2_ fold changes of common ISGs at 8, 16, 24 hours of INF induction.

**Table S4.** Oligos used in the study.

**Table S5.** Gene lists and gene expression changes in mimics experiments.

